# Dis-integrating the fly: A mutational perspective on phenotypic integration and covariation

**DOI:** 10.1101/023333

**Authors:** Annat Haber, Ian Dworkin

## Abstract

The structure of environmentally induced phenotypic covariation can influence the effective strength and magnitude of natural selection. Yet our understanding of the factors that contribute to and influence the evolutionary lability of such covariation is poor. Most studies have either examined environmental variation without accounting for covariation, or examined phenotypic and genetic covariation without distinguishing the environmental component. In this study we examined the effect of mutational perturbations on different properties of environmental covariation, as well as mean shape. We use strains of *Drosophila melanogaster* bearing well-characterized mutations known to influence wing shape, as well as naturally-derived strains, all reared under carefully-controlled conditions and with the same genetic background. We find that mean shape changes more freely than the covariance structure, and that different properties of the covariance matrix change independently from each other. The perturbations affect matrix orientation more than they affect matrix eccentricity or total variance. Yet, mutational effects on matrix orientation do not cluster according to the developmental pathway that they target. These results suggest that it might be useful to consider a more general concept of ‘decanalization’, involving all aspects of variation and covariation.

## Introduction

The evolutionary response to natural selection requires phenotypic variation. As such, the mechanisms generating phenotypic variation and covariation (**P**) are of fundamental importance. Most studies focus on the genetic component in **P**, summarized as the genetic covariance matrix **G**, due to its role in the response to natural selection as summarized in the multivariate version of the breeder's equation (Lande 1979; Lande and Arnold 1983). Thus, the generation of heritable components of variation, governed by mutation and recombination, is broadly studied. However, non-heritable sources of phenotypic (co)variation, often summarized through **E**, remain important for our understanding of both the magnitude and direction of response to natural selection through its contributions to **P** (i.e., **P**=**G**+**E**). The basic measure of the strength of selection - the selection differential - depends on **P** via the covariance between the phenotype and fitness (the Robertson-Price identity; Robertson 1966; Price 1970). In addition to its influence on the magnitude of selection, **E** influences the response to selection via **P** as summarized in Lande's (1979) equation (∆Z=**GP**^−1^*s*). Empirical evidence shows that E is shaped by genetically-defined reaction norms to differences in environment, and can thus be adaptive and amenable to directional and stabilizing selection (Hill and Mulder 2010). In a multivariate phenotype, the covariances in **E** affect the orientation of the response to selection in addition to its magnitude. Even in cases when **E** does not facilitate adaptive evolution directly, it can certainly impede it and possibly bias its trajectory towards a secondary peak (Burger 1986). Yet, our understanding of how this environmental covariation is generated during development is incomplete, as is our understanding of how it evolves.

Mutational and environmental perturbations have often been utilized to examine how the environmental component of phenotypic variation changes, particularly in the context of canalization (Waddington 1952; Dworkin 2005a,b; Levy and Siegal 2008; Hallgrímsson et al. 2009; Hill and Mulder 2010; Paaby and Rockman 2014; and references therein). A number of these studies have found an increase in phenotypic variance in response to perturbations, often attributed to decanalization, and due in part to a release of cryptic genetic variation (Dworkin 2005a,b; Paaby and Rockman 2014; and references therein). Yet, both increases and decreases of environmental variance have also been observed without the release of genetic variation (Debat et al. 2006; Dworkin and Gibson 2006; Debat et al. 2009; Hallgrímsson et al. 2009). Most studies, however, have examined the effect of mutational perturbations on the variance of a trait or on the total variance across several traits (e.g., Dworkin 2005a; Levy and Siegal 2008), whereas in multivariate systems other aspects of covariation need to be considered.

Covariance matrices are commonly characterized based on their size, shape, and orientation (Jones et al. 2003; Hohenlohe and Arnold 2008), as illustrated in Fig. 1. Matrix size is often measured as the total variance in this context (sum of the individual trait variances) and it quantifies variation overall, irrespective of how it is distributed among different directions (Fig. 1A; henceforth referred to as total variance). The orientation of the matrix refers to the direction in which the primary axes of covariation are pointing (Fig. 1B), and is determined by the relative contribution of each trait to each eigenvector. Thus, a change in matrix orientation reflects a change in the direction in which most of the variation is concentrated, and therefore a change in the combination of traits that are more easily attainable through evolution. The eccentricity of the matrix describes its shape – how much it deviates from a hypersphere – and it quantifies how evenly the variation is distributed among the different directions (Fig. 1C), irrespective of orientation and total variance. A more eccentric matrix (i.e. more cigar-shaped) means that there is considerably more variation along the first axis relative to other axes, and therefore a fewer trait combinations that are easily attainable. To the extent that the observed covariation reflects integrating factors, higher eccentricity reflects a more integrated body plan.

**Figure 1.**
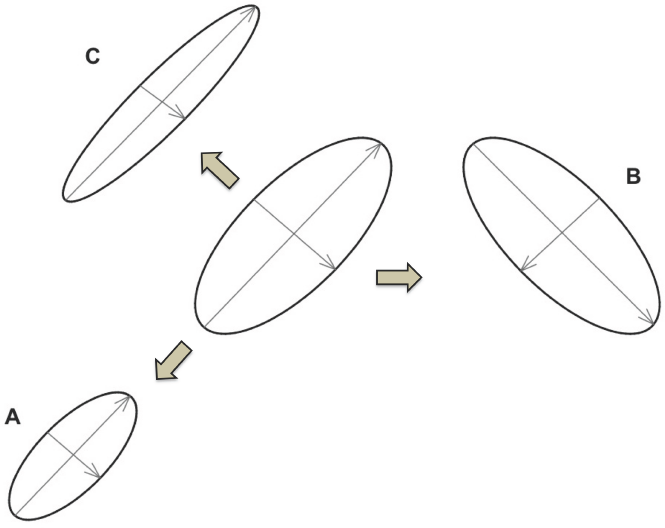
Illustration of the different matrix properties. Ellipses represent 95% confidence interval of a normal bivariate distribution, and arrows (ellipse axes) represent orthogonal axes of covariaton. Matrices are presented as bivariate for illustration, but the principles are the same for multivariate. Ellipses A, B, and C represent a change in only one property each, relative to the ellipse in the middle. A, lower total variance; B, perpendicular orientation; C, higher eccentricity.

Previous studies provide mixed expectations as to how the different properties of the covariance matrix might change relative to each other and relative to mean shape. Moreover, the association between changes to matrix orientation and eccentricity has mostly been studied so far for **P** and **G** rather than **E**. Simulations by Jones et al. (2003, 2004, 2007, 2012) and Revell (2007) suggest that the orientation of **G** is more likely to change than eccentricity under most conditions, including effects due to population size, magnitude, and orientation of mutational correlation and stabilizing selection, and different modes of directional selection. In addition, higher eccentricity enhances the stability of the orientation under most conditions. The magnitude and stability of eccentricity and total variance, on the other hand, are mostly influenced by population size rather than mutational correlations and selection. Empirical studies have found evidence that both eccentricity and orientation can be either stable or labile across populations and species at different phylogenetic scales (Arnold et al. 2008; de Oliveira et al. 2009; Porto et al. 2009; Haber 2015a,b). Unfortunately it is not clear to what extent the expectations from these studies (examining **G** and **P**) are relevant to **E**. Changes in **G** reflect a combination of influences from mutation and other evolutionary forces. The structure of **E**, on the other hand, reflects the interaction between the environment and development, and different genotypes differ in their sensitivity to the environment. Thus, while strong directional selection can deplete genetic covariation (Homrigh et al. 2007, Walsh and Blows 2009), altering matrix properties of **G**, it is not yet clear how this should alter the matrix properties of **E** as well.

Theory suggests that strong selection, both disruptive and directional, can lead to an increase in the total variance of **E** (Hill and Mulder 2010). Hallgrímsson et al. (2009) postulated that when the total variance increases due to developmental perturbations, matrix eccentricity could either increase or decrease, depending on how the perturbed developmental process contributes to the original patterns of covariation. Eccentricity would increase with total variance if the targeted pathway plays a major role in integrating the measured traits, thus adding variance to the major axis of covariation. When the perturbed pathway does not contribute much to the major axis of covariation, changes in variance are distributed among the smaller subsequent eigenvalues, thus reducing eccentricity. Indeed, empirical studies have found both increases and decreases of eccentricity with an increase in the total variance of **E** (Hallgrímsson et al. 2009). In addition, studies have found mutational effects on matrix orientation of **E** that could not be readily associated with developmental pathways or mutational effect size (Hallgrímsson et al. 2006; Debat et al. 2009; Jamniczky and Hallgrímsson 2009). Most studies, however, did not include a direct comparison of the different properties of **E** across a wide range of mutations, nor how these properties change relative to mean shape. Thus, we are left with little understanding of the lability of **E**.

*Drosophila* wing shape is an ideal system to address these questions, and in particular how mutational perturbations influence the different aspects of the covariance structure of **E**. The development of the wing of *D. melanogaster* has been a model system for over 70 years, and it is one of the best genetically characterized model systems. The extensive set of genetic tools, mutant lines, and ability to generate many genetically identical individuals reared under common environments enables the study of mutational effects on **E**. Variation in both size and shape of the *Drosophila* wing has a long history of research, and sophisticated high dimensional representations of shape can be generated in a high throughput manner (Houle et al. 2003; Pitchers et al. 2013). Numerous studies have shown that mutational perturbations can have profound effects on mean shape (Weber et al. 2005, Breuker et al. 2006; Debat et al. 2006; Dworkin and Gibson 2006; Debat et al. 2009, Debat et al. 2011). The influence of mutational perturbations on variance has been more mixed, with some studies seeing no effect, some find a general increase (i.e., de-canalization; Debat et al. 2009, Debat et al. 2011), and others finding both increase and decrease (Debat et al. 2006; Dworkin and Gibson 2006). Some evidence for the effect of mutational and environmental perturbations on matrix orientation has been shown in these studies as well. However, changes to covariance and shape have not been compared directly, and other aspects of the covariance structure, such as eccentricity, have not been included before.

To better understand the evolutionary lability of the **E** matrix, we used a set of induced mutations measured against a common co-isogenic wild type (Dworkin and Gibson 2006; Debat et al. 2009), as well as a panel of strains derived from natural populations, and examined the relationship between mean shape and different aspects of the covariance. Thus, all strains were reared under the same carefully-controlled conditions, and all mutant lines had the same genetic background, minimizing unknown sources of variation. We observe that mean shape varies more freely than the covariance structure, and that matrix orientation varies more than – and independently from – its eccentricity and total variance.

## Material and Methods

### Drosophila strains

Insertional mutations (caused by the insertion of P-elements, marked with *w*^*+*^) in genes involved with the TGF-β and EGFR signaling pathways were provided from the Bloomington Stock Center (Table 1). These represent a subset of the alleles that were first described in Dworkin and Gibson (2006) and were chosen because of their important roles in the growth, patterning, and shape determination of the *Drosophila* wing. All insertions were initially introgressed into the wild-type lab strain Samarkand (Sam) for at least 10 generations. The Samarkand genotype was marked with *w*^−^ (white eyes) so flies with insertions could be distinguished by a rescue of the eye color phenotype. Introgressions were performed by repeated backcrossing of females bearing the insertion to males of Sam. Prior to generating flies for the experiments described in this study, the alleles were backcrossed to Sam for an additional four generations to remove any *de novo* mutations that had accumulated in those lines (relative to Sam) since the original introgression procedure. Selection was based entirely on the presence of the eye color marker, precluding unwitting selection for wing phenotypes. All crosses were performed using standard cornmeal-molasses media, in a 24° C Percival incubator on a 12/12-hr light/dark cycle with 60% relative humidity (Dworkin and Gibson 2006). The mutant strains used here are part of a larger unpublished study (Dworkin), and were chosen in part based on sample size (>50/genotype).

**Table 1.**
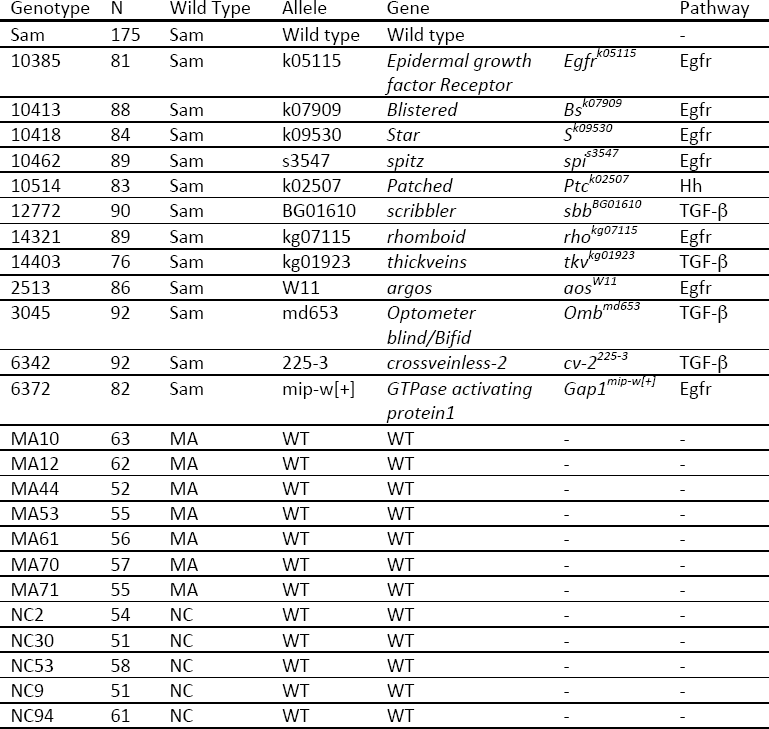
Information about the genotypes included in this study.

### Experimental setup

After the additional four generations of backcrossing back to Samarkand, crosses between each mutation and Samarkand were set up in vials, allowing females to lay for 2-3 days, so that egg density was low (generally less than 60 flies/vial). The temperature of the incubator was maintained at 24° C, and monitored carefully for fluctuations, and vial position was randomized within the incubator on a daily basis to reduce any possible edge effects. As larvae crawled out of the media, a piece of paper towel was added to each vial to provide additional pupation space. After eclosion and sclerotization flies were separated, on the basis of eye color, into individuals without the P-element-induced mutations (w−) and heterozygotes for the P-element, and then stored in 70% ethanol.

### Strains derived from natural populations from North Carolina and Maine

In order to broaden the comparative basis we included a set of naturally-derived Iso-female lines of *D. melanogaster*. Since too little is known about the scale of differences we might expect for matrix properties, even for fruit flies, it helps to look at the range of differences found among natural populations. Since shape has no natural zero (it is on interval scale), we chose Sam as the point of comparison for all strains, mutants as well as naturally-derived.

The naturally-derived lines were established separately in the summer of 2004 at a Peach orchard in West End, North Carolina (NC2) and in a blueberry field in Cherryfield, Maine (courtesy of Marty Kreitman). To generate inbred lines, full-sib mating was performed for 15–20 generations (Goering et al 2009; Reed et al 2010). Flies were reared at 25C (+/− 1°C) in a 12/12 light/dark cycle at 50% humidity. We note that a different incubator was used for these lines (compared to the mutant lines), but preliminary analyses for several of the P-element mutant lines demonstrated highly consistent results with respect to mean shape (not shown). Seven Maine and five NC2 lines, each with at least 51 males, were used for this study.

### Data collection

A single wing from each fly was dissected and mounted in 70% glycerol (~20 wings per replicate vial on average, >50 wings per genotype, males only; see Table 1). Wings were imaged at 40X magnification on an Olympus DP30BW camera mounted on an Olympus BX51 microscope using ‘DP controller’ V3.1.1 software. All images were saved as greyscale tiff files. To extract landmark and semi-landmark data, we followed a modified protocol for the use of the WINGMACHINE software (Houle et al. 2003) as detailed in Pitchers et al. (2013). First, we used “tpsDig2” (Rohlf 2010) software to record the coordinates of the two starting landmarks needed by WINGMACHINE. We then used WINGMACHINE to automatically fit nine B-splines to the veins and margins of the wing (Fig. 2A). We reviewed each splined image and manually adjusted the control points as necessary. The x and y coordinates of 14 landmarks and 34 semi-landmarks (Fig. 2A) were extracted using additional software (CPR) developed by Marquez and Houle (2014). The data were checked for visual outliers on scatter plots, and putative outliers were examined, and either fixed in WINGMACHINE or deleted. The resulting dataset was then passed on to **R** for further analysis.

**Figure 2.**
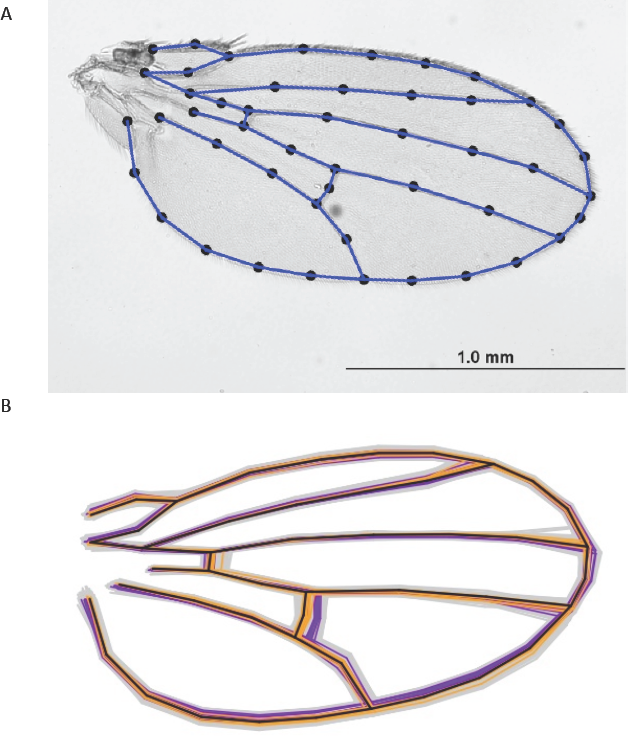
**A,** A representative wing image with fitted splines and the location of landmarks and semilandmarks along them. **B,** Individual variation in the sample (grey) relative to Sam (black), mutant means (orange), and means for naturally-derived lines (purple).

### Preparation of data

All specimens from all genotypes were superimposed together using Generalized Procrustes Analysis and projected into tangent space (function *gpagen* in the **R** package *geomorph 2.1.1;* Adams and Otarola-Castillo 2013). The semi-landmarks were optimized using minimum Procrustes distance. This resulted in a space of 96 Procrustes coordinates with 58 degrees of freedom (Zelditch et al. 2004). Replicate effect and the allometric effect of size were removed from the shape coordinates by fitting a linear model, excluding their interaction, carried out separately for each genotype. The predicted mean configuration was then added back to the residuals to maintain the differences among genotypic means.

After regression, the dataset was passed on to the LORY program for calculating the interpolated Jacobian-based data (Marquez et al. 2012). LORY uses spatial interpolation to evaluate shape deformation at predetermined evaluation points throughout the wing. It starts by creating a tessellated grid, with the landmarks and semi-landmarks as the vertices (Fig. 3A). The centroids of the resulting triangles are considered the evaluation points. Given an interpolation function, it then calculates the Jacobian matrix that describes the shape deformation at each point (e.g., Fig. 3B). The log-transformed determinant of the Jacobian matrix quantifies the amount of expansion or contraction at each evaluation point relative to the mean configuration. Thus, the interpolation takes into account information from the whole configuration of landmarks, as well as from the rest of the sample through the mean configuration, while quantifying local shape changes. The Procrustes coordinates are mathematically transformed from a single multidimensional trait into a multivariate set of traits. Just like interlandmark distances (and unlike the Procrustes shape variables), each of the LORY variables can be interpreted independently. The whole dataset can thus be analyzed using conventional multivariate techniques (e.g., PCA, multivariate regression), yielding results that are more easily interpretable as well (Marquez et al. 2012).

**Figure 3.**
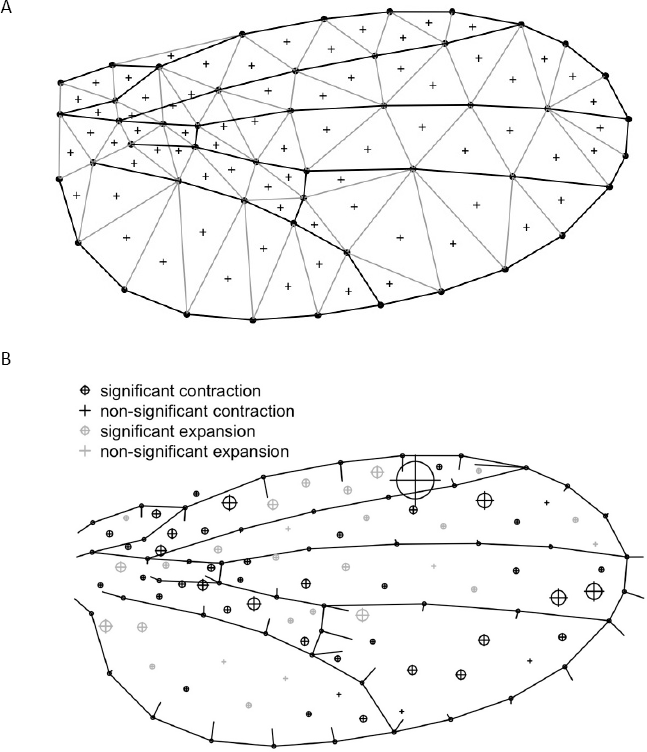
Illustrating the LORY method for interpolating shape changes from Procrustes data of genotype *Bs*^*k07909*^. **A**, the tessellation scheme for choosing evaluation points. The landmarks and semi-landmarks (dots) are used as vertices. The centroids of the resulting triangles (crosses) are the final evaluation points. **B**, shape differences between the genotype mean and Sam inferred by calculating the Jacobians around each evaluation point, scaled by 100 (crosses and crossed circles). Bars represent the direction and change of differences between Sam and the sample mean at the landmarks and semi-landmarks based on the Procrustes data, scaled by 10X.

The interpolation function allows the researchers to incorporate and test an explicit model for the distribution of shape changes across the configurations, rather than relying solely on the Procrustes superimposition. The covariance matrix from the Procrustes residuals is modified based on that explicit model. The most common interpolation function used in geometric morphometrics studies is the Thin Plate Spline (TPS), which assumes the object is rigid and therefore penalizes local deformations more than global ones. In this study we compared TPS with another interpolation function implemented in LORY, the Elastic Body Spline (EBS), which assumes the object is elastic and penalizes local deformations less than global ones (Marquez et al. 2012). Therefore, like the Procrustes superimposition, TPS tends to spread the variation more globally than EBS. Because TPS and EBS make different assumptions about the distribution of variance among landmarks, they also make different assumptions about the integration pattern of the wing. EBS largely implies less integration and more compartmentalization. For this study system, we consider EBS to be a more suitable model *α priori*, because the development of the fruit fly wing has been shown by most studies to proceed in a relatively compartmentalized manner in which variation tends to be locally contained (Zecca and Struhl 2002; Barrio and de Celis 2004; Cook et al. 2004; Martín and Morata 2006; Ziv et al. 2012; cf. Klingenberg and Zaklan 2002 and Debat et al 2003).

Each of the three dataset – the Procrustes coordinates and the LORY variables based on TPS and EBS – was reduced to its first 30 dimensions using Principal Component Analysis, covering 99% of the variation in the data (Fig. S1). All subsequent analyses, including estimation of the covariance matrices, are based on these PCA scores. Thus, we have three sets for each of these measures: mean shape distances, total variance, covariance distances (i.e., differences in orientation), and eccentricity. However, results based on Procrustes data were highly correlated with the Jacobian-based data, using both TPS and EBS (Figs. S2–S3). Therefore, we chose to present below only results based on EBS, and provide results for the other two datasets in the supplemental material. Thus, for *Drosophila* wing shape, using the LORY variables did not make a substantial difference (compared to simply using Procrustes residuals). However, this could change for other cases, depending on the specific covariance structure of any given set of configurations (see examples in Marquez et al. 2012).

### Quantification of variation and covariation

Mean shape distances were calculated as the Euclidean distance for both the interpolated data and the Procrustes-based data. Since the Procrustes data were projected to tangent space after superimposition (see above), this is equivalent to Procrustes distance. Variance-covariance (VCV) matrices were quantified based on their total variance, shape, and orientation (see Fig. 1). Total variance is the trace of the matrix (i.e., sum of its eigenvalues). Matrix shape was characterized based on its eccentricity (Jones et al. 2003), and calculated as the ratio of the largest eigenvalue to the total variance, which is the inverse of Kirkpatrick (2009)'s effective number of dimensions (Kirkpatrick 2009):

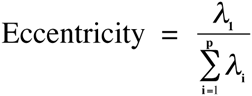

Where **λ**_1_ is the largest eigenvalue, **λ**_i_ are the remaining i^th^ eigenvalues, and *p* is the number of variables in the VCV matrix. In addition, we calculated eccentricity as the relative standard deviation of the eigenvalues (rSDE), also known as integration level (Van Valen 1974; Pavlicev et al. 2009a; Haber 2011). As developed originally by Van Valen (1974) for covariance matrices, rSDE is also scaled by the total variance, thus measuring matrix shape only (Haber 2011, 2016). However, rSDE yielded essentially the same results as the above measure of eccentricity (Figs. S4 and S13), and will not be considered here further.

Similarity in matrix orientation was quantified using Random Skewers (Cheverud et al. 1983; Cheverud and Marroig 2007; Marroig et al. 2011), which measures the average similarity between two matrices in their response to random unit vectors. We used 5000 random vectors drawn from a normal distribution with mean 0 and variance 1, normed to unit length (Marroig et al. 2011). A distance metric was then calculated as √(1-*r*^2^), where *r* is the Random Skewers value for that pair of matrices, and normalized using Fisher's *z*-transformation (Jamniczky and Hallgrímsson 2009). In accordance with previous studies, this metric is referred to here as “covariance distance”. Two other distance metrics were calculated as well: a modification of Krzanowski's (1979) metric following Zelditch et al. (2006), and the relative eigenvalues metric of Mitteroecker and Bookstein (2009). However, these two methods proved unstable for our dataset under resampling, yielding unreliable confidence intervals. At the same time, their estimated values followed the same pattern as Random Skewers (see online supplementary material). Moreover, Random Skewers is the only method of the three that is related to evolutionary theory (Hansen and Houle 2008). Therefore, we present below only results based on Random Skewers.

Confidence intervals for shape distances, total variance, and eccentricity, were estimated using a non-parametric bootstrap procedure with a BCa correction (DiCiccio and Efron 1996; Carpenter and Bithell 2000) using 999 iterations. The BCa correction was necessary because the pseudovalue distribution is expected to be biased upward when the statistic is bounded by zero, and to depend on its mean when bounded by both zero and one. These confidence intervals were used for evaluating significant differences as well. Confidence intervals for covariance distance were estimated using a Jackknife procedure. Each Jackknife pseudovalue was calculated by leaving out one specimen in one of the two samples that are being compared. The confidence interval was calculated as the 95% of the Jackknife distribution, without a bias correction. The Jackknife was preferred in this case because the bootstrap (both parametric and non-parametric) consistently resulted in distributions that were highly biased upward, often excluding the observed value, invalidating the BCa correction. This is a commonly found phenomenon in similar studies and has not yet been properly addressed in the literature to the best of our knowledge. Here we found that Jackknife without a bias correction provided fairly symmetric distributions around the observed values with reasonably wide confidence intervals, whereas the bias-corrected jackknife yielded extremely narrow intervals that often excluded their observed value (i.e., they were “over-corrected”).

In order to further compare covariance changes with shape changes we used Principal Coordinates Analysis to generate a shape space and a covariance space, based on all pairwise distances. In each of these spaces, genotypes are located based on how different they are from each other in either shape (i.e., shape space) or matrix orientation (i.e., covariance space). The two spaces where then superimposed together using a symmetric Procrustes superimposition including scaling (function *protest* in **R** package *vegan 2.0–10*; Oksansen et al. 2013). The superimposition ensures that the two spaces are comparable in terms of both magnitude and direction of change, just as it does with landmark configurations (Peres-Neto and Jackson 2001). The sum of squared deviations between the two spaces after superimposition provides a measure of correlation between them. In addition, we calculated the disparity of mutants and naturally-derived strains - for shape and covariance - as the sum of variances of their respective scores in the joint space, including all 20 PCoA axes. Although the configurations are scaled by their respective centroid size during the superimposition, the protest function then rescales the rotated configuration (the covariance space in this case) proportionally to the target configuration (the shape space in this case) so that their size (total variance) in the joint space is comparable but not necessarily the same. This allows us to evaluate the disparity of one space relative to the other.

We used MANOVA (Pillai's Λ) to test the effect of the developmental pathway on the mutants' PCoA scores. As mentioned above, each genotype involved one mutation that targeted either the TGF-β or the EGFR signaling pathway (see Table 1). This analysis allowed us to test whether mutations clustered based on the pathway they targeted.

In order to estimate the effect of the high dimensionality of our data on sampling variance, we repeated all analyses using a reduced dataset, for which we kept only the 12 landmarks and omitted all semi-landmarks. This dataset had 20 degree of freedom (12*2–4), which is less than half our smallest sample size (45). This reduced dataset yielded very similar results as the full dataset, for all properties, in terms of both the observed values and the confidence intervals (Figs. S5–S7). The fact that the confidence intervals are similar is especially relevant here, indicating directly that sampling variance is not affected by reducing the dataset dimensionality from 56 to 20.

All analyses were carried out in **R** 3.2.4. All scripts and data will be made available on DRYAD and on the Dworkin lab github repository.

## Results

### Both mean shape and matrix orientation vary considerably among mutant and naturally-derived strains

All mutants differ significantly in their mean shape from the Sam wild type (Table 2 and Fig. 4; see also Fig S8), as determined by the lack of overlap between their confidence intervals and the benchmark of zero distance from Sam (horizontal dashed line in Fig. 4). Most mutant strains are more similar to the Sam wild type than any of the naturally-derived strains. Yet, two of the mutants (genotypes 3045 and 10413, genes *Omb*^*md653*^ and Bs^k07909^ respectively) show magnitudes of shape change as extreme as any of the naturally derived strains, suggesting that the range of mutations included here likely provide a reasonable representation of what we might observe in nature. The orientation of the covariance matrix of all mutant strains also differ significantly from Sam (Table 2 and Fig. 5): none of the covariance distances include zero in their confidence interval. Unlike mean shape, however, most of the mutant strains differ from Sam just as much as the naturally-derived strains do.

**Figure 4.**
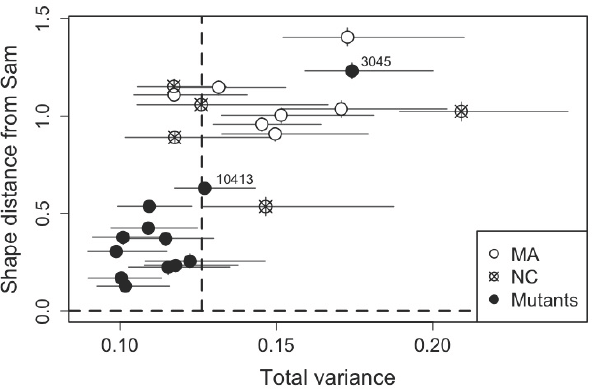
Mean-shape distance from Sam plotted against the total variance within each strain. Bars are 95% confidence intervals. Dashed lines represent values for Sam. Labeled mutants are mentioned in the text.

**Table 2.**
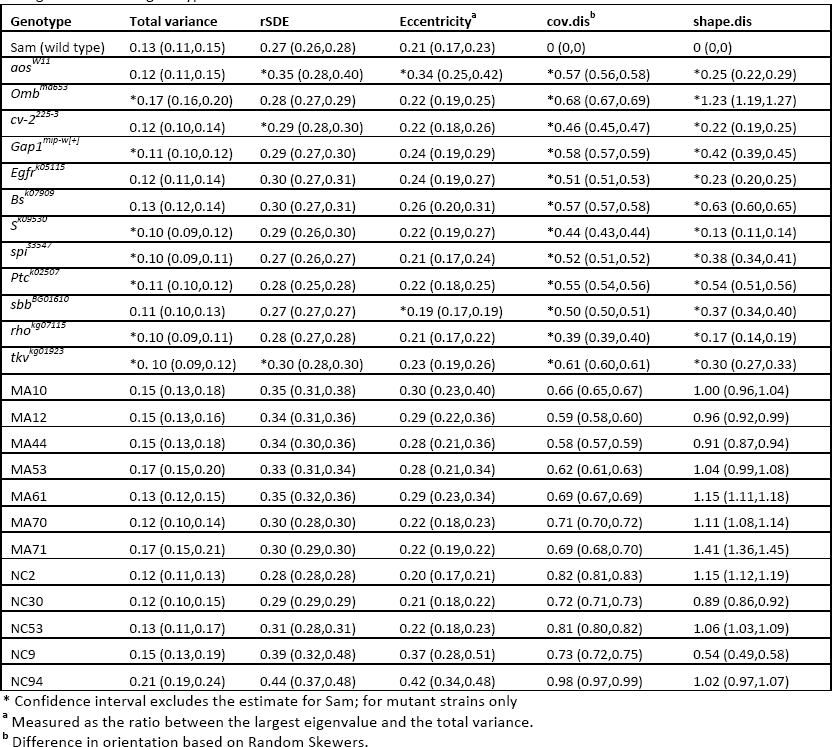
Matrix properties of all genotypes, and their distance from Sam in terms of matrix orientation (cov.dis) and mean shape (shape.dis). All measures are based on Jacobians, using EBS, and are therefore proportional to the mean configuration of that genotype.

**Figure 5.**
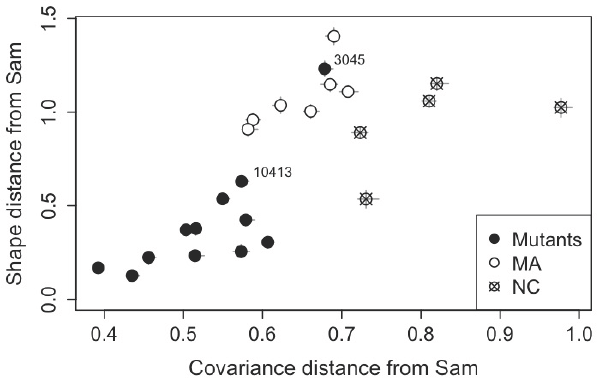
Mean-shape distance plotted against covariance distance (i.e, difference in matrix orientation; estimated by Random Skewers), relative to Sam. Bars are 95% confidence intervals. Labeled mutants are mentioned in the text.

### Several mutant strains have lower environmental variance than their wild type

Most mutations resulted in a lower total variance than their co-isogenic Sam wild-type (Table 2 and Fig. 4). Six of the 12 mutant strains exclude the estimate for Sam from their confidence interval (Table 2), thus indicating a significant difference. Two of the significant strains (*Ptc*^*k02507*^, *tkv*^*kg01923*^) are part of the TGF-β pathway, and the other four (*Gap1*^*mip-w[+]*^, *S*^*k09530*^, *rho*^*kg07115*^, *spi*^*s3547*^) contribute to EGFR signaling (Table 1). The only strain that has substantially and significantly higher total variance is *Omb*^*md653*^, which also differs greatly in its mean shape. In contrast, the total variance within each of the naturally-derived strains is mostly higher than the total variance within Sam. In addition, there is a positive linear relationship between the total variance within the mutant strains and their shape distance from Sam (Fig. 4).

### A positive non-linear association between changes in mean shape and matrix orientation

There is a weak but positive association between covariance distance (i.e., difference in matrix orientation) and shape distance (Fig. 5), relative to Sam. This relationship, however, is not linear (Table 3), suggesting that the covariance distance is somewhat bounded at the upper range. These findings are further supported by comparing the shape and covariance PCoA spaces (Fig. 6, see also Figs S9–S12). These spaces are based on all pairwise comparisons, rather than comparing to Sam only, and were superimposed for comparability. The superimposition scales the two spaces to a common scale, centers, and rotates them so that they are comparable in terms of both magnitude and direction of change. The grey solid lines indicate the deviation between the two spaces. The sum of squared deviations between the two spaces is 0.669, and the coefficient of determination is small yet significantly different from zero (R^2^ = 0.33; p≤0.001 with 999 permutations). Thus, the two spaces are substantially different from each other, but not unrelated. The first two PCoA axes cover 58% of the variation in the joint space (Fig. 6), while the third and forth axes cover only 10% and 6% of the variation, respectively. The first PCo1 axis mostly separates the naturally-derived strains from the mutants and Sam, for both shape and covariance. The disparity of shape is more than twice as large as that of covariance (i.e., orientation differences) (0.042 and 0.020 respectively; calculated over all 20 PCoA dimensions that resulted in eigenvalues larger than zero). The disparity of both shape and covariance is larger for the naturally-derived strains than for the mutants (Fig.6A). The mutants do not cluster by pathway (Fig. 6B), for either shape (Pillai's Λ=0.44, p=0.24) or covariance distance (Pillai's Λ=0.34, p=0.52).

**Table 3.** Non-linear regression of shape distances on covariance distances (relative to Sam)

|  | Estimates | t value | P value |
| --- | --- | --- | --- |
| Intercept | -2.83<br>(-4.75, -0.92) | -3.07 | 0.0057 ** |
| Linear term | 9.07<br>( 3.20, 14.94) | 3.21 | 0.0042 ** |
| Non-linear term | -5.19<br>(-9.57, -0.81) | -2.47 | 0.022 * |

Model: $Y = X + X^2$ ; Y is shape distance; X is covariance distance
Tested sequentially so that the non-linear term is added to the linear term
Residual standard error: 0.25 on 21 degrees of freedom
Multiple R-squared: 0.63, Adjusted R-squared: 0.60
F-statistic: 18.29 on 2 and 21 DF, p-value: 2.52e-05

**Figure 6.**
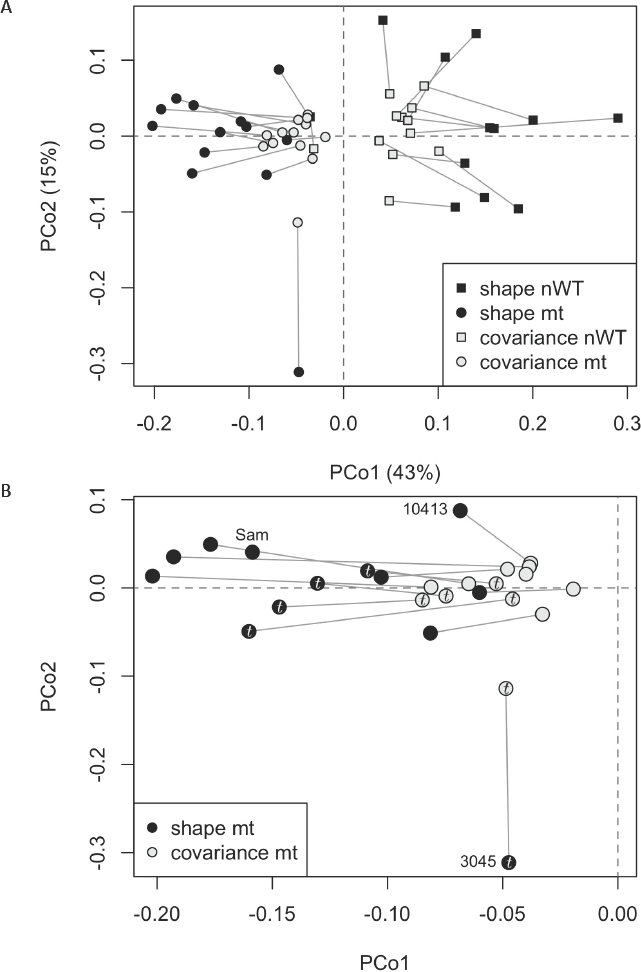
The first two axes of the covariance space superimposed onto shape space. Both spaces are based on all pairwise distances. Solid grey lines indicate the deviation between the two spaces for a given genotype. **A**, All genotypes; **B**, Mutants only, including Sam. Mutants that affect the TGF-β pathway are marked by *t.* All other mutants affect the EGFR pathway. nWT, naturally-derived wild types; mt, mutants. Labeled mutants are mentioned in the text.

### Most mutations do not influence matrix eccentricity, despite effects on mean shape, total variance, and matrix orientation

Most of the mutations do not have a substantial effect on the eccentricity of covariation (i.e., matrix shape), and include the estimate for Sam well within their confidence interval (Table 2). In addition, there is no clear relationship between eccentricity and total variance (Fig. 7A). The only mutations that caused a significant change in eccentricity are *aos*^*W11*^, which increases relative to Sam, and *sbb*^*BG01610*^, which decreases relative to Sam (genotypes 2513 and 12772, respectively). These mutations do not differ much from Sam for total variance and mean shape (Fig. 7 and Table 2). *Omb*^*md653*^, on the other hand, differs significantly and substantially from Sam in its total variance and shape (see Table 2), and yet is very similar in its eccentricity to Sam. Most of the naturally-derived strains have a higher eccentricity than Sam (Fig. 7A), showing a greater difference from Sam than the mutants do. Similarly, there is no clear association between eccentricity and shape distance from Sam (figure 7B), or between eccentricity and covariance distance (i.e., orientation differences) from Sam (Fig. 7C). The same picture emerges when eccentricity is calculated using the relative standard deviation of the eigenvalues (rSDE; Table 2 and Fig. S13), and based on Procrustes and TPS data (Figs. S14–S15).

**Figure 7.**
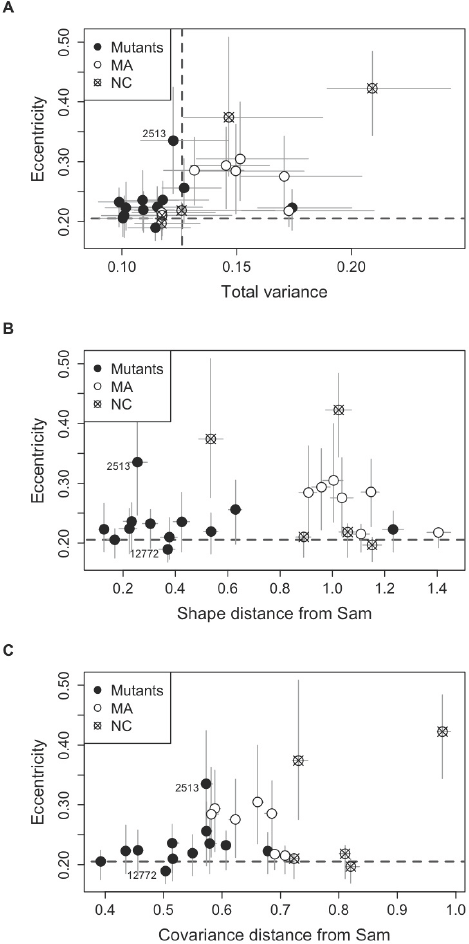
The association between eccentricity and (**A**) total variance within each genotype, (**B**) their mean-shape distance from Sam, and (**C**) their covariance distance from Sam. Dashed line represents the eccentricity of Sam. The apparent positive relationship between eccentricity and total variance is no greater than expected by chance based on parametric simulations. Labeled mutants are mentioned in the text.

## Discussion

The pattern and magnitude of phenotypic covariation remains central to selection theory (Robertson 1966; Price 1970; Lande and Arnold 1983). Although it cannot usually facilitate adaptation by itself, the environmental component of phenotypic covariation – **E** – can impede and divert the population response to selection (Burger 1986). Despite the importance of this for understanding evolution, factors that influence the lability of **E** are poorly understood. For univariate measures, environmental and genetic stressors have been shown to increase variance in **E** in numerous situations (Dworkin 2005a; Hallgrímsson et al. 2009; Paaby and Rockman 2014; and references therein), but little is known about how such perturbations influence other properties of integration and covariation.

Using the wing of *Drosophila melanogaster* as a model system, we found that mean shape and matrix orientation vary substantially among the co-isogenic mutants, as well as the naturally-derived strains (Table 1 and Fig. 5). By superimposing the covariance space onto shape space we were able to compare their distribution in terms of both magnitude and direction of change and show that shape has changed to a greater extent than matrix orientation (Fig. 6). In contrast, mutations had little influence on the total variance and eccentricity of the matrix. Whereas total variance is positively associated with mean shape, it is not associated with eccentricity, nor is eccentricity associated with matrix orientation. Together, these findings suggest that the potential of the covariance structure to change is more limited than that of mean shape, and that different properties of the covariance structure can change independently from each other. Thus, even though these mutations greatly affect the location of genotypes in morphospace (i.e., mean shape), the availability of different trait combinations to natural selection (matrix orientation) is not affected as much. Moreover, the total amount of variation (matrix size) and its distribution among the different trait combinations (eccentricity) are not affected much either. It is worth noting that some of the environmental effects (such as rearing temperature and density) were highly controlled in this study, both within and among strains. Thus, our experimental design may represent a lower bound for the environmental contributions to covariation, and the patterns observed are likely an underestimate of the possible effect sizes.

The common expectation is that mutational perturbations would cause a disruption to the developmental system, leading to a higher variance and greater shape changes (Dworkin 2005b; Hall et al 2007; Hallgrímsson et al. 2009; Hill and Mulder 2010; and references therein). Our findings are consistent with the common expectation in that greater decanalization is indeed associated with larger changes to mean shape, but surprising in the sense that most mutants have a lower variance than their co-isogenic wild type (albeit significant for only 6/12 genotypes), and thus seem to be more canalized. If we had used linear measurements, we would have expected to find a positive correlation between the mean and the variance merely due to scale. However, such a technical relationship is not expected for Procrustes shape data because all specimens are scaled by their centroid size during the superimposition. Thus, the positive association we find likely reflects more than a technical relationship. In addition, the difference in variance between strains is not likely to be due to differential survival, because the individuals within each strain are genetically identical and viability was high enough for all genotypes. Thus, the decrease in total variance within the mutants likely reflects a higher canalization of wing shape. Dworkin and Gibson (2006) have also observed such mixed results for mutant strains, using a smaller set of landmarks. A related study (Debat et al 2009) showed a more consistent increase in total variance, even for some of the same mutations. However, the genetic background of the strains in Debat et. al (2009) was Oregon-R (rather than Samarkand), which has been shown to be more sensitive to mutational perturbations under some conditions (Chari & Dworkin 2013).

Increases in the environmental component of phenotypic variance have been explained in previous studies by a nonlinear relationship between the trait mean and its underlying developmental parameters (Klingenberg and Nijhout 1999; Hallgrímsson et al. 2009). The nonlinearity arises from the expectation that the variation expressed around the mean phenotype can vary for different phenotypes, reflecting different levels of canalization. A steeper slope around a given phenotype implies a greater amount of variation, and therefore less canalization, associated with that phenotype. It is often assumed that when development is perturbed the mean would shift to a less canalized section on the curve and express greater variation. It is possible, however, that the mean would shift into more canalized regions, rather than less canalized, resulting in a decrease of variance rather than an increase (Hill and Mulder 2010). In other words, Sam is more sensitive to micro-environmental influences than the mutants and less sensitive than the natural strains, at least in our study. It is also possible to have system-wide canalizing factors (such as chaperon proteins; Rutherford and Lindquist 1998) operating simultaneously, masking the effects of local perturbations and stabilizing the variance (Hallgrímsson et al. 2006). Or, the mutations could increase redundancy and complexity thus increasing stability (Hallgrímsson et al. 2006; Levy and Siegal 2008). All of these possible explanations would be consistent also with the positive association we found between mean shape changes and total variance.

A possible explanation for the higher variance within the natural strains (relative to Sam) could be related to the time and amount of inbreeding among the strains (Whitlock & Fowler 1999; Whitlock et al. 2002). Inbreeding in *Drosophila* does represent a form of “genetic stress” as deleterious recessive alleles are made homozygous (and their effects are phenotypically expressed), often leading to an increase in environmental variance. We would therefore expect the natural strains to be de-canalized as well. However, the Sam wild-type has also undergone extensive inbreeding, and is near isogenic (Chandler et al. 2014). Thus, a satisfactory explanation for this particular observation eludes us.

In accord with previous studies (Dworkin and Gibson 2006; Debat et al. 2009; Jamniczky and Hallgrímsson 2009), covariance matrices of the mutant strains do not cluster according to the developmental pathway that their underlying mutations presumably affect. This could point to the complexity of the developmental system, further supporting the palimpsest model suggested by Hallgrímsson et al. (2009). It could also mean that these pathways do not contribute to the structure of **E** as effectively and consistently as previously postulated. A detailed analysis of the covariance pattern is needed to further investigate the extent to which the modularity structure of the fruit fly wing follows *a priori* developmental models, and how it is disrupted by developmental perturbations.

It is difficult to compare differences in eccentricity to differences in orientation directly because of their different dimensionality and somewhat different method for calculating the confidence intervals. However, mutant eccentricity varies from that of Sam within a smaller portion of its possible range (0.2–0.43; 23%) compared to how much their orientation varies from Sam (0.37–0.77; 40% without the Fisher's *z*-transformation). These findings are in accord with simulations by Jones et al. (2003, 2004, 2012) for **G**, suggesting that eccentricity is likely to vary less than orientation for a given population size under most combinations of genetic architecture, selection regimes, and phylogenetic scale. It is also consistent with empirical evidence for **P** from Haber (2015a,b), showing that matrix orientation has varied greatly among closely-related ruminant species, whereas eccentricity has remained largely the same throughout most of bovid history, and has only varied within 33% of its possible range among other ruminants.

As with many studies of covariation, sampling variance could have a substantial impact on our results, considering the high dimensionality of the data, and especially on estimates of covariance. Several studies (i.e., Hill and Thompson 1978; Meyer and Kirkpatric 2008; Pavlicev et all. 2009b) have shown that the leading eigenvector tends to be overestimated due to sampling variance, and the trailing eigenvalues underestimated. This would affect eccentricity the most. Simulations carried by Haber (2011) indicate that eccentricity is only biased (overestimated) for low values (rSDE<0.2) with sample sizes lower than 45 and number of variables 35 or lower. Our sample sizes are all higher than 45, and most are higher than 60, with 56 degrees of freedom, so largely equivalent to those simulations, and all of our rSDE values are higher than 0.2. There is no reason to expect other measures of eccentricity to be affected differently as they are all very tightly correlated and reflect the same property (the distribution of the eigenvalues). In addition, the power analysis in Haber (2011) indicates that increasing sample size above 40 adds little to the statistical power of rSDE. Therefore, it is reasonable to assume that our eccentricity estimates are largely unbiased by sampling variance. With regards to other properties, mean shape and total variance are scalars, known to be relatively robust to sampling (Zelditch et al 2004). In contrast, matrix orientation is probably the most sensitive property, but it is impossible to say how sampling variance would affect it as it depends largely on the covariance structure itself. However, since sample size is similar for all mutant lines, its effect should be about the same as well. Moreover, repeating the analyses with a reduced dataset of 12 landmarks (see methods), yielded very similar results as the full dataset for all properties, in terms of both the point estimates and their confidence intervals (Figs. S5–S7). The fact that the confidence intervals are similar is informative in this context as it indicates that the sampling variance is not substantially affected by reducing the dataset from 56 to 20. Pitchers et al. (2014) also found that the number of families (the relevant sample size for that study) had relatively modest effect on various measures of matrix properties of **G**.

To conclude, previous studies have shown an increase in variance to be clearly associated with decanalization. In this study, however, we find that mutants altered the orientation of the covariance matrix more often than its total variance. Mutations do not only affect trait means and variances, but aspects of covariances as well, though this has not been part of the general formulation so far. Thus, our study suggests that it might be useful to consider a more general concept of decanalization.

## Acknowledgements

We thank Ellen Larsen, Jeff Conner, Will Pitchers, Vincent Debat, Adrien Perrard, Handling Editor Ruth Shaw, Associate Editor Ricardo Azevedo, and two anonymous reviewers for comments that improved clarity of this manuscript. This material is based in part on work supported by the National Science Foundation under Cooperative Agreement No. DBI-0939454. Any opinions, findings and conclusions or recommendations expressed in this material are those of the author(s) and do not necessarily reflect the views of the National Science Foundation. This work was supported by NIH grant 1R01GM094423-01 to ID. Funding for AH was provided by the Council of Higher Education, Israel, and BEACON Center for the Study of Evolution in Action at Michigan State University.

